# Phosphorylation of Ser81 in human AGT reversibly inactivate enzyme function and mimics catalytic defects of certain PH1-causing mutations

**DOI:** 10.1101/2025.04.14.648764

**Authors:** Sara Milosevic, Mario Cano-Muñoz, Eduardo Salido, Noel Mesa-Torres, Angel L. Pey

**Affiliations:** Hosptial Universitario de Canarias, Tenerife, España; Universidad de Granada, Granada, España

**Author notes:** Corrpespondence to.

## Abstract

Phosphorylation is fundamental to modulate protein function and stabilty. There have detected about 300000 site-specic Phosphorylation sites in over 20000 human proteins. However, only 5% of the sites have been experimenrally characterized. In this work, we investigated a phosphorylation event in AGT, an important enzyme due to its detoxifying role of glyoxylate and hundreds of mutations cause a rare disease (primary hyperoxaluria type I or PH1). We analyzed the effect of phosphomimetic mutations Ser81 on the WT proten, the common polymorpshim LM, and the most common disease-associted variants (LM-G170R and LM-I244T). Using biochemical, biophysical and cell biology approaches, we show that phosphorylation at S81 dramatically affects PLP/PMP binding pose and disrupts enzyme activity, without pertubing its subcellular location to peroxisomes. This reversible phenotype is similar to the irreversible effects of some PH1-causing mutaions. Thus, we provide evidence for a novel regulatory mechanism for PH1, in health and disease.

## 1. Introduction

Proteins undergo co- or post-ransalatios modificications (named as PTMs) that modulate their function, regulation and stability [1–15]. These events have critical implications in health and disease. According to the Phosposite Plus ® database [16](accessed the 13^th^ April 2025), there are ~0.5 million PTMs described in over 20000 human proteins by high-throughput methods. Only a 5.3 % have been characterized in detail. It is noteworthy that ~60% of the PTMs regard phosphorylation (of Ser, Thr and Tyr residues). Overall, ~36% of all PTMs are related with phosphorylation of Ser residues. What is the relevance of Ser phosphorylation in health and disease? We show here that the effects can be monumental.

We have used as a model of conformational disease using human alanine:glyoxale 1 (named as AGT). Alterations in the function of these enzyme leads to a rare disease called primary hyperoxaluria type I (PH1) [17–19]. However, the role of site-specific phosphorylation on AGT function and stability has almost not being investigated [15]. In human population, the WT sequence exists in 80 % of the alleles, while 20% contains several genetic variants (relevantally, two mutations, p.P11L and p.I340M) [20,21]. Out of six potential phosphorylation sites, four involve Tyr modifications (Figure 1), which requires no conventional mutational phosphomimetic strategies [22].

**Figure 1.**
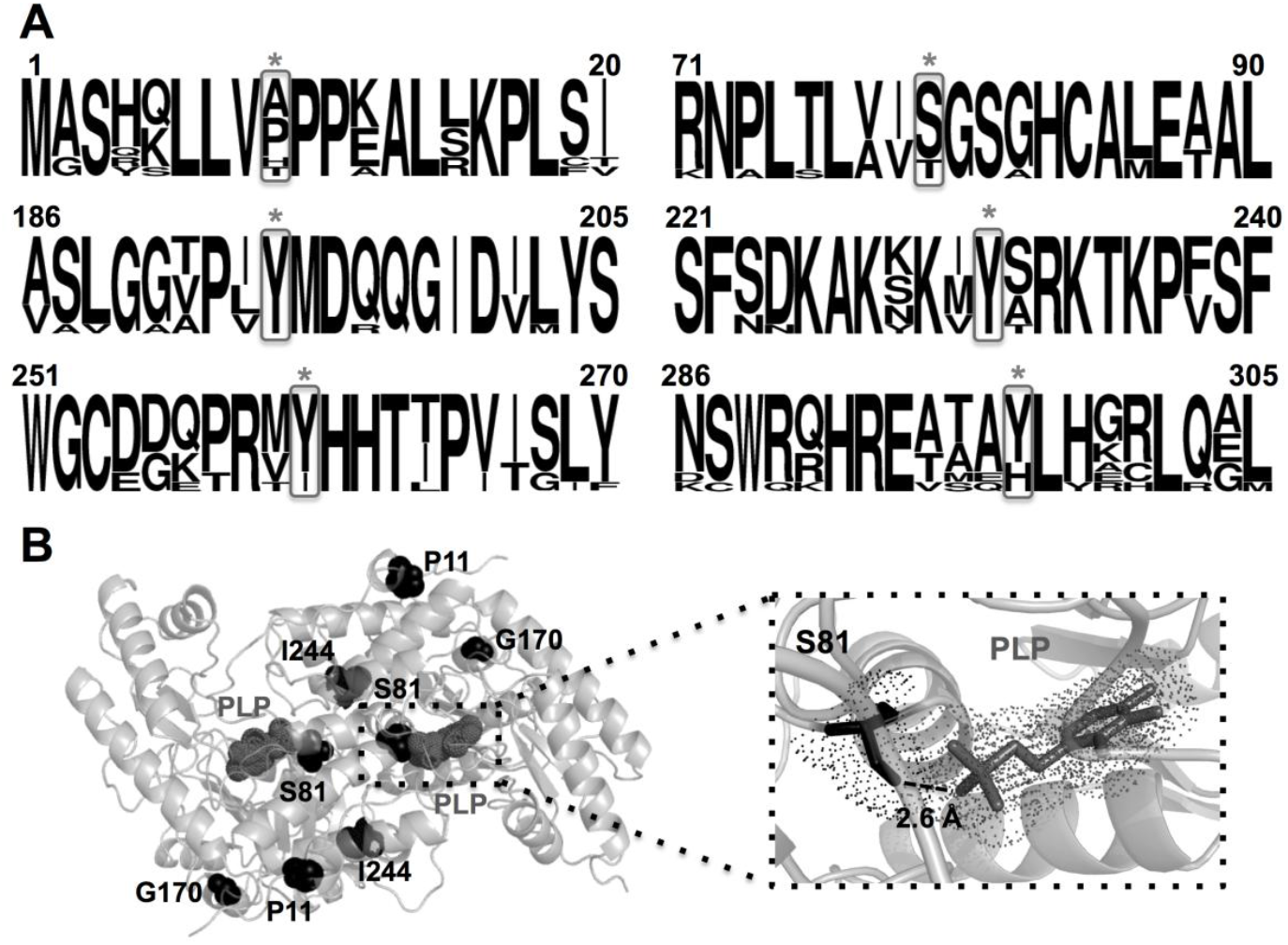
Selection of S81 as phosphorylation site to be studies in AGT variants associated with Primary hyperoxaluria type I. A) Sequence-alignment statistical analyses of AGT sequences from mammals for the vicinity of the five phosphorylation sites described in Phosphosite Plus ® (from top to the bottom: T9, S81, Y194, Y231, Y260 and Y297). Sequences included were those from human, orangutan (*Pongo abelii*), monkey (*Macaca mulatta*), baboon (*Papio anubis*), dog (*Canis lupus familiaris*), horse (*Equus caballus*), cat (*Felis catus*), cow (*Bos taurus*), rabbit (*Oryctolagus cuniculus*), rat (*Rattus norvegicus*) and mouse (*Mus musculus*) enzymes. The figure was produced using WebLogo (https://weblogo.berkeley.edu/logo.cgi). B) Structure of human AGT dimer (PDB 1H0C) showing the location of P11, G170, I244, S81 and the PLP molecule. The lower panel shows a close view of the PLP binding site with S81 at very close distance (as short as 2.6 Å between the side chain O of S81 and one of the phosphate oxygens in PLP).

We show here that phosphorylation of Ser 81 have dramatic effects on enzyme activity and cofactor binding. These effects are not phenotype-dependent. Our work thus provides a new layer to understand the physiology and pathology of AGT.

## 2. Materials and Methods

### 2.1. Protein expression and purification

Mutations to p.S81A and p.S81D were introduced into pCOLDII plasmids containing AGT WT, LM, LM-p.G170R or LM-p.I244T [23] using standard mutagenesis protocols and confirmed by DNA sequencing of the entire cDNA. BL21(DE3) *E. coli* strains were transformed with these plasmids, and protein expression was carried out by induction of these bacterial cultures using 0.4 mM IPTG for 4-6 h at 4°C. AGT proteins were purified from soluble extracts by metal affinity chromatography and subsequent size-exclusion chromatography as described in [23,24]. Apo-proteins were produced by treatment with L-Ala and mild acidic pH as previously described [24]. Proteins were stored in Na-HEPES 20 mM NaCl 200 mM pH 7.4 and their concentration was measured spectrophotometrically using ε_280_= 47000 M^−1^·cm^−1^ for the AGT monomer.

### 2.2. Overall transamination activity measurements

AGT specific activity was measured at 37ºC typically using 2.5 to 4 µg·ml^−1^ purified AGT protein (up to 50 µg·ml^−1^ for variants containing the mutation p.S81D). The reaction mixture contained 10 mM glyoxylate, 100 mM L-alanine, 150 µM PLP in 100 mM K-phosphate pH 8. The reaction time was set to 2 min and the reaction quenced by using trichloroacetic acid (25% w/v final concentration). Samples were clarified by centrifugation at 21000 g for 10 min at 4ºC and supernatants were stored at −20ºC. Pyruvate formed in these reactions was determined using lactate dehydrogenase and NADH 0.2 mM at 37ºC. The reaction was monitored as the change in A_340 nm_ in Tris-HCl 1 M pH 8 using quartz cuvettes in a Agilent 8453 spectrophotometer at 25°C. These amounts of pyruvate were determined using calibration curves of of pyruvate under the same conditions. For each variant we carried out at least four different replicas, and the results were presented as mean±s.d.

### 2.3. Spectroscopic and light scattering measurements

All spectroscopic measurements were carried out in Na-HEPES 20 mM NaCl 200 mM pH 7.4 unless otherwise indicated. UV-visible absorption spectra were collected in an Agilent 8453 diode-array spectrophotometer at 25°C using 1 cm path length cuvettes and 20 μM AGT proteins as purified. To trigger the formation of PMP, a final concentration of 0.2 M L-alanine was added and the reaction was allowed to proceed for at least 10 min at 25°C before spectra collection. To determine whether the PMP formed was released upon addition of L-alanine, these reactions were filtered using 30 kDa cut-off concentrators (VIVASPIN® 6, Sartorius), and the spectra of filtrates was acquired under similar conditions to those used with AGT proteins. Circular dichroism (CD) measurements were performed at 25 °C in a Jasco J-710 spectropolarimeter equipped with a Peltier temperature controller. AGT samples for CD measurements in the Near UV-Visible range were prepared as described above for absorption measurements. Spectra were recorded at a scan rate of 100 nm·min^−1^ with a 2 nm bandwidth and five scans were registered and averaged, using 5 mm path-length cuvettes and appropriate blanks without protein were registered and subtracted. For Far-UV CD measurements, AGT proteins were prepared at 5 μM in K-phosphate 20 mM pH 7.4, and spectra were recorded at a scan rate of 100 nm·min^−1^ with a 1 nm bandwidth and six scans were registered and averaged, using 1 mm path-length cuvettes and appropriate blanks without protein were registered and subtracted. Absorption and CD spectra were recorded from two different preparations of each AGT variants (two independent purifications) and averaged. Dynamic light scattering (DLS) was measured using a DynaPro MSX instrument (Wyatt) in a 1.5 mm path-length cuvette and using 10 μM AGT proteins as purified. Twenty-five spectra were acquired at 25°C for each DLS analysis, averaged and used to determine the hydrodynamic radius assuming spherical scattering particles (i.e. the Stokes-Einstein approach). Radii were expressed as mean ± s.d. from 4-6 replicates.

PLP binding to apo-proteins was measured under pseudo first-order kinetic conditions. PLP (5-100 μM) were always in a large excess compared to apo-AGT (typically 0.4 μM in monomer). Protein samples were thermostatized in 1 cm quartz-cuvettes at 25°C for five minutes, and PLP was added to an appropriate final concentration, and mixed manually (the dead time was 10-40 s, registered and appropriately considered in the calculations). Time-dependent emission fluorescence was acquired in a Cary Eclypse spectrofluorimeter (Varian), using an excitation wavelength of 280 nm and an emission wavelength of 340 nm (5 nm slits) at 25°C. Kinetic traces were fitted to a single exponential function to provide the observed rate constant *k*_obs_. Under pseudo first-order conditions, this rate constant is *ideally* related to the association equilibrium constant K_a_, the second-order association rate constant *k*_on_ and the first-order dissociation rate constant *k*_off_ as follows:

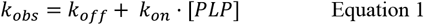

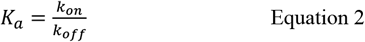

where [PLP] is the total PLP concentration. Thus, a linear fit of equations 1-2 to the *k*_obs_ vs. [PLP] data provides the values of the apparent equilibrium and kinetic constants for PLP binding, as well as their corresponding fitting errors. Dissociation constants (K_d_) were simply calculated considering that these are equal to inverse of K_a_ (=1/K_a_).

### 2.4. Thermal stability

Thermal stability of AGT proteins (1 μM in monomer) was determined by following the changes in fluorescence emission intensity (exc. 280 nm, em. 360 nm; slits 5 nm) using a Cary Eclipse spectrofluorimeter (Varian). Holo-AGT contained proteins as purified with 100 μM PLP, while apo-proteins were prepared from apo-protein stored solutions. All protein solutions were prepared in Na-HEPES 20 mM NaCl 200 mM pH 7.4 and placed in 1 cm path-length cuvettes. After 5 min of incubation at 20°C, thermal scans were carried out from this temperatura up to 70-95°C at a scan rate of 2°C·min^−1^. Upon subtraction of pre- and post-transition linear baselines, the apparent denaturation temperature (*T*_m_) was determined as the temperature at which half of the denaturation signal was achieved. Values reported are mean±s.d. from 3-4 replicas for each protein variant.

### 2.5. Molecular modeling

In contrast to holo-AGT, no crystal structure is available for the PMP-AGT state. Therefore, we generated a PMP bound form of the enzyme following previous reports [25,26]. We used the structure of holo-AGT (PDB ID: 1H0C)[27] to place PMP in the active site reproducing the PLP binding mode by manual docking and maintaining water molecules present in the original crystallographic structure. Structure relaxation performed by energy minimization following previous works [25,26]. All heavy atoms were fixed to allow added hydrogens to adjust in the newly generated environment. Then, PMP ligand and side-chain of every residue within a sphere of 20 Å were free to move during the simulation.

Then, YASARA molecular modeling software was used to generate the single mutants p.S81A and p.S81D [28,29]. To remove bumps and correct the covalent geometry, the structure was energy-minimized with the NOVA force field [30], using a 8 Å force cutoff and the Particle Mesh Ewald algorithm [31,32] to treat long-range electrostatic interactions. After removal of conformational stress by a short steepest descent minimization, the procedure continued by simulated annealing (timestep 2 fs, atom velocities scaled down by 0.9 every 10^th^ step) until convergence was reached, i.e. the energy improved by less than 0.05 kJ/mol per atom during 200 steps.

2D schematic representation of the binding interactions between PMP and the enzyme’s binding site were created with PoseView [33]. The H-bonds are shown as black dashed lines, and the hydrophobic contacts are represented by residue labels and spline segments along the contacting hydrophobic ligand parts.

### 2.6. Experiments in CHO cells

Chinese hamster ovary (CHO) cells (ATTC, USA) were grown in alpha-minimal essential medium (α-MEM, Lonza, Germany) supplemented with glutamine, penicillin/streptomycin and 5% fetal bovine serum, on 13 mm glass coverslips placed in 6-well plates (Avantor, Spain). Cell transfections were performed using AGT cDNA variants subcloned in pCIneo plasmids (Promega, USA) with FectoCHO reagent (Sartorius, Germany), following manufacturer’s guidelines.

The following AGT variants were tested: S81 in both major (WT) and minor (LM) alleles, and D81 in both major (WT) and minor (LM) alleles.

After 24 h, cells were fixed with buffered formaline (Avantor, Spain) for 10 min., permeabilized with 0.1 triton-X and immunofluorescence was performed as previously described [23]. Cellslls were labeled with two different sets of antibodies: a) guinea pig anti-AGT (1:1000) and rabbit anti-PMP70 (1:1000) to assess colocalization of AGT and the peroxisomal marker; and b) guinea pig anti-AGT (1:1000) and rabbit anti-mitochondria (1:1000) to assess colocalization of AGT and the mitochondria. After 1 hr incubation with primary antibodies, coverslips were washed with phosphate buffered saline (PBS) and incubated for 30 min. with the secondary antibody mix (Alexa Fluor-488 goat anti-guinea pig IgG and Alexa Fluor-555 goat anti-rabbit IgG). After PBS wash, coverslips were mounted on glass slides with PBS-glycerol containing Hoesch-33342 to counterstain the nuclei. Digital images were acquired with a fluorescence microscope (Zeiss, Germany) using a 60x objective with immersion oil and sequential blue-green-red channel emmission filters. Image analysis was performed with Fiji version of ImageJ software (NIH).

## 3. Results

### 3.1. p.S81D affects PLP binding and thermal stability with little effect on the overall dimeric conformation

Twelve AGT variants, AGT WT, LM, LM-p.G170R and LM-p.I244T (set S81) containing the mutation p.S81A (set A81) or p.S81D (set D81) were expressed in and purified from *E*.*coli* soluble extracts. As isolated, these purified variants showed similar overall secondary structure (Figure 2A-B and Figure S1A) and similarly behaved as dimers in solution (Figure 2C and Figure S1B), indicating that the p.S81D and p.S81A mutations hardly affected he overall structure of the AGT dimer.

**Figure 2.**
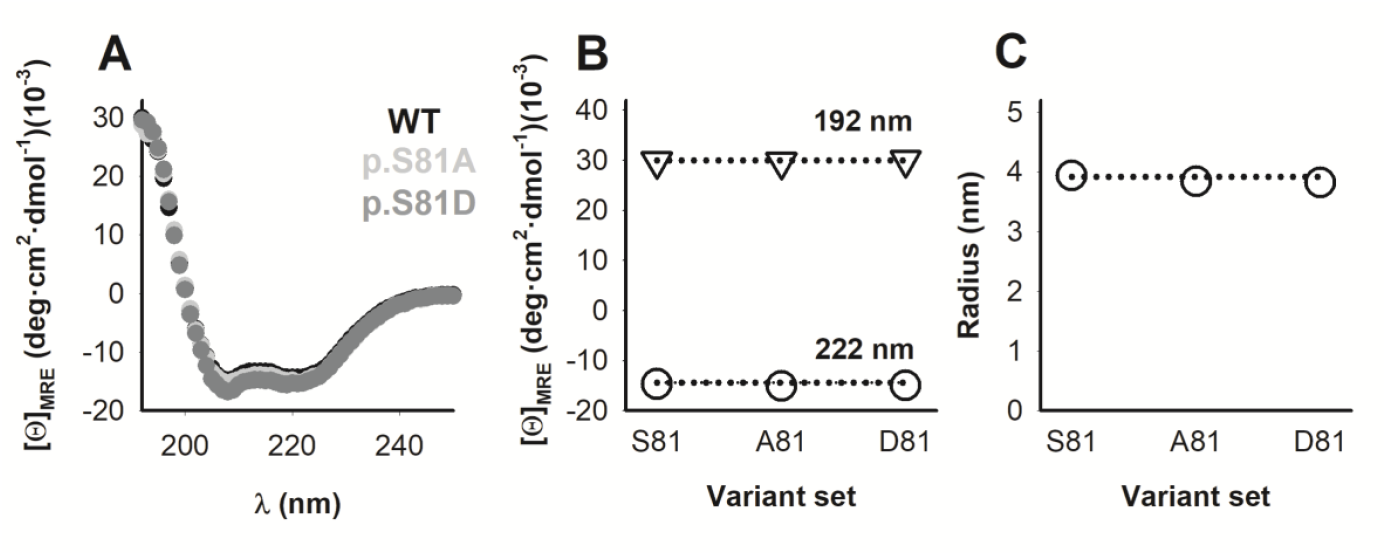
Overall conformation of AGT proteins. A-B) Far-UV CD spectra for AGT proteins as purified (A). For each variant, spectra are the average from two different purifications. In B, we show data for the mean residue ellipticies at 192 nm and 222 nm (average±s.d. for four AGT variants belonging to the S81, A81 and D81 sets, respectively). C) Hydrodynamic radius for AGT proteins as purified from DLS analyses. Data are the average ± s.d. from 4-6 independent measurements. In B and C, the horizontal dotted lines indicate values for AGT WT as reference and errors are smaller than the symbols.

Binding of PLP to apo-AGT variants often causes a dramatic increase in the thermal (conformational) stability of the protein, increasing the T_m_ by about 25°C [23,24,26,34]. In addition, the LM variant showed lower stability than the WT protein, and this destabilization was enhanced by the p.G170R and p.I244T mutations primarily in the apo-state [23,35]. Strikingly, when we carried out thermal denaturation experiments, the behavior of the D81 set clearly differed from those of the S81 and D81 sets (Figure S2). Indeed, when we compared those variants D81 with those containing either S81 or A81, we observed significant diferences. The introduction of p.S81D led to significant stabilization of apo-AGT, with an average increase of 7.7±1.2°C of variants containing p.S81D vs. those containing S81 (Figure S2). In addition, an excess of PLP did not stabilize those variants containing p.S81D, those leading to a much lower thermal stability than those containing S81 (ΔT_m_= −23.1±2.4°C)(Figure S2). The effects of the p.S81A mutation were much milder, with ΔT_m_= −1.5±0.5°C and 2.4±1.3°C, for holo- and apo-AGT when these were compared with controls containing S81 (Figure S2). Consequently, the perturbation introduced by p.S81D seems to stabilize the apo-AGT in all genetic backgrounds but prevents stabilization of AGT upon PLP binding.

Spectroscopic analyses of the PLP bound to AGT variants provided additional insight evidence for function alterations in the D81 set. AGT shows very high levels of covalently bound PLP forming a Schiff base with K209 [23,25,27]. An unaffected binding pose of PLP can be identified from the spectral features in the absorption as well as its CD spectra (Figure 3A-C and Figure S3A-B). However, those variants containing p.S81D showed alterations in the electronic properties and local microenvironment of PLP. The maxima of the spectra for the S81 and A81 sets was found to be 428±3 nm (average of eight variants), while the presence of p.S81D blue-shifted these maxima by about 35 nm (mean±s.d of 392±1 nm for these four variants). In addition, CD spectra also suggested severely distorted binding of PLP to variants containing p.S81D, since the dichroic signal associated with bound PLP was absent for these mutants (Figure S3B).

**Figure 3.**
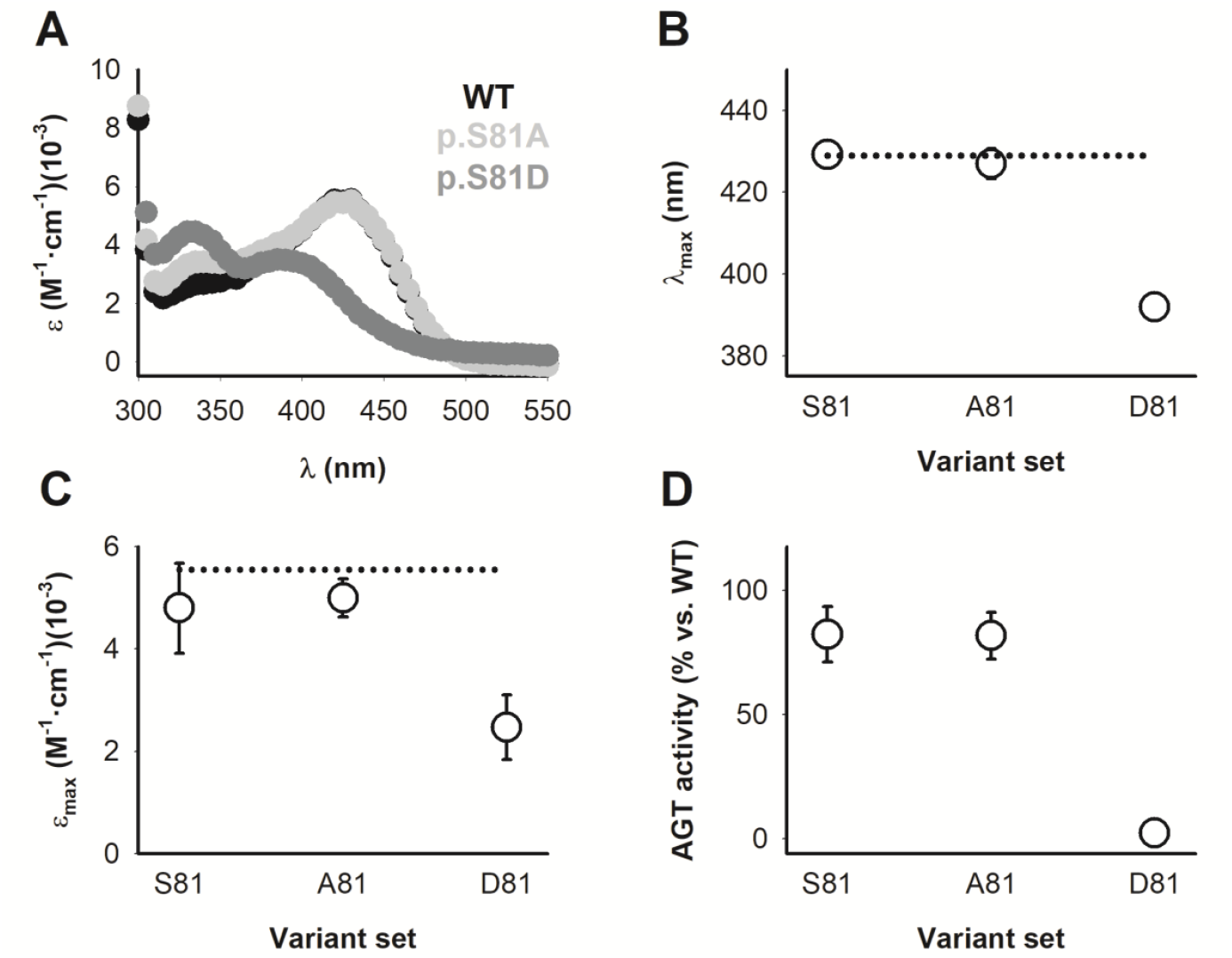
Spectroscopic analyses support that the mutation p.S82D distorts PLP bound conformation and abolishes AGT activity. A) Near UV-Visible absorption spectra of AGT WT, p.S81A and p.S81D variants. B-C) Values of the maxima of absorption specta (B) and the extinction coefficient at the maximum (C) for Holo samples. D) Overall transamination activity. Data are mean±s.d. of four different variants (WT, LM, LM-p.G170R and LM-p.I244T) for the S81, A81 and D81 sets. Each variant was normalized considering the value of WT AGT. Horizontal dashed lines in panes B and C correspond to the values of WT AGT.

### 3.2. p.S81D inactivates AGT and prevents the conversion of PLP into PMP in the presence of L-alanine

To assess whether variants containing p.S81D may affect the overall transaminase AGT activity, we carried out activity measurements with high concentrations of L-ala and glyoxylate as substrates and in the presence of an excess of PLP (Figure 3D and S1C). Importantly, we observed that variants forming the S81 and A81 sets showed similar activity (1.39±0.17 mmol Pyr·h^−1^·mg^−1^ protein) while the mutation p.S82D essentially abolished it (activity of 0.04±0.05 mmol Pyr·h^−1^·mg^−1^ protein for the D81 set).

To address whether this distorted PLP binding pose in the D81 set could prevent the formation of PMP in the transamination reaction, we incubated AGT samples with a large excess of L-alanine and evaluated the formation of PMP by changes in absorption and CD spectra. Again, we observed that variants forming the S81 and A81 sets showed virtually full conversion of PLP into PMP, as reflected by the dissapearance of the strong absorption band at 428±3 nm and the appearence of a strong band with a maximum at 329±2 nm with a higher extinction coefficient (from 4.9±0.7 to 7.3±0.9 mM^−1^·cm^−1^ at the maximum; all values are mean±s.d. for all S81 and D81 variants) associated with PMP formation (Figure 4A-B and S3A)[25]. However, these changes were much smaller for the D81 set, with a small red-shift in the maximum (from 392±1 nm to 406±2 nm) and a small increase in the extinction coefficient at the maximum (from 2.5±0.6 to 3.2±1.1 mM^−1^·cm^−1^) (Figure 4A-B). Similarly, the dichroic signal associated with PMP formation is not observed in D81 variants (Figure 4C and Figure S3B). Thus, the lack of transaminase activity in these variants is presumably due to their inability to catalyze the amine transfer between the bound substrate and the PLP cofacto

**Figure 4.**
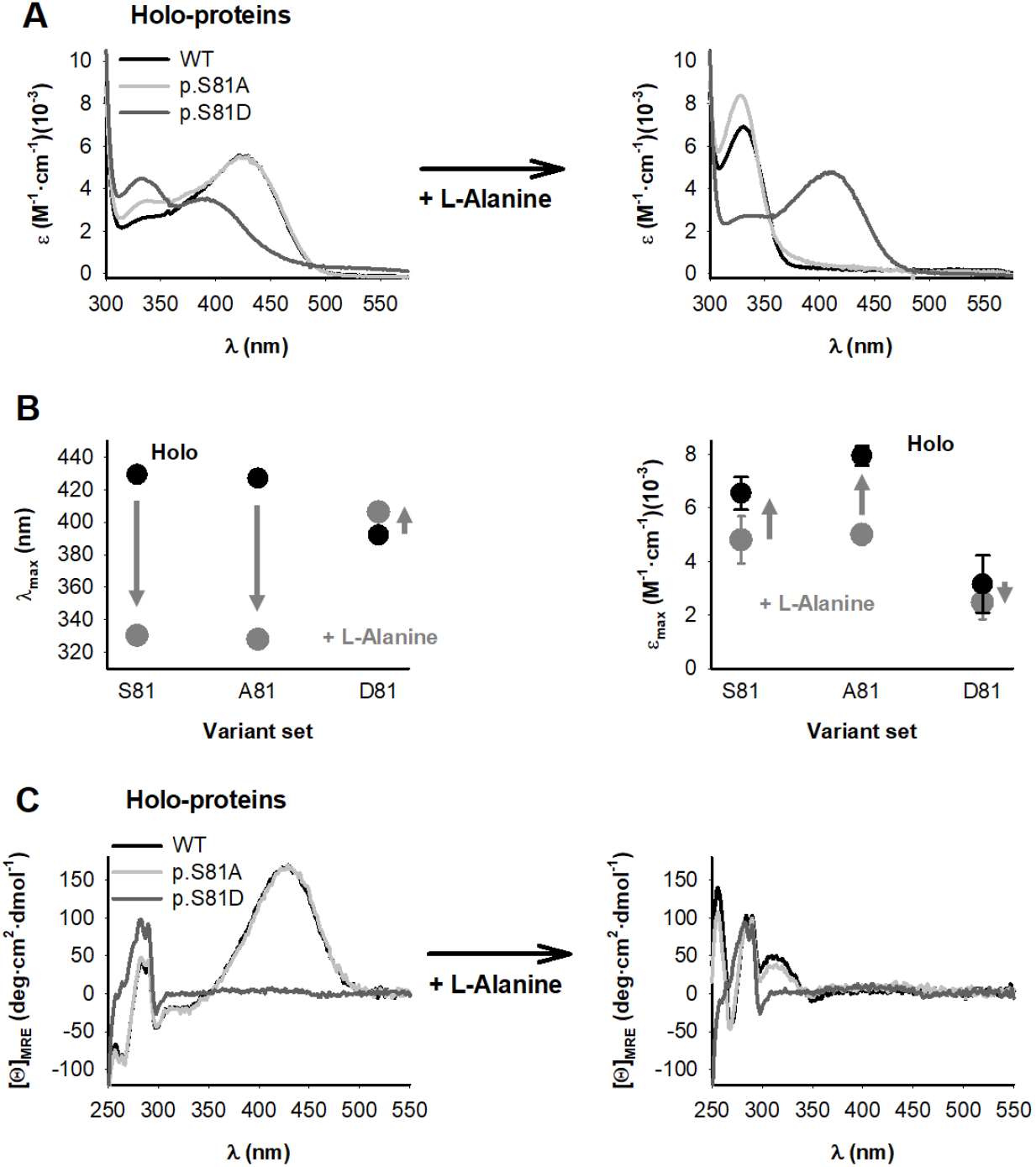
Spectroscopic analyses support that the mutation p.S82D prevents PLP to PMP transamination in the presence of L-Alanine. A) Near UV-Visible absorption spectra of AGT WT, p.S81A and p.S81D variants before (left panel) and after (right panel) addition of 200 mM L-Alanine. B) Values of the maxima of absorption specta (left panel) and the extinction coefficient at the maximum (right panel) for Holo samples before and after addition of 200 mM L-Alanine. Data are mean±s.d. of four different variants (WT, LM, LM-p.G170R and LM-p.I244T) for the S81, A81 and D81 sets. Vertical arrows indicate the change observed upon addition of L-alanine. C) Near UV-Visible CD spectra of AGT WT, p.S81A and p.S81D variants before (left panel) and after (right panel) addition of 200 mM L-Alanine. Spectra were corrected for dilution and are the average from two experiments using two different protein preparations for each variant.

### 3.3. The mutation p.S82D severely reduced PLP binding affinity while the mutation p.S82A may reduced PMP binding affinity

So far, we have found that the p.S81D mutation severely affects the PLP binding pose and likely facilitates the release of PLP from its complex with AGT. From a simple perspective, alterations in PLP binding energetics and kinetics may arise from effects of this mutation on the complex with the protein as well as in the protein in the unbound state. The former should primarily affect the kinetics of PLP dissociation from the complex (*k*_off_) while the latter should affect mostly the association rate constants (*k*_on_). We must note that, although these interpretations are simple, intuitive and very reasonable (thus, appealing), the possibility of more complex scenarios for PLP binding cannot be excluded (i.e. the existence of different free and bound states, see [36]

Consequently, we determined both rate constants as well as estimated the equilibrium dissociation constant (*K*_d_) for all variants from PLP binding kinetic experiments (Figure 5A-B). PLP binding to the S81 variants is clearly much slower that those of S81 and A81 set (k_on_ values are 6.1±1.2 for the D81 set vs 84±21 M^−1^·s^−1^ for all S81 and A81 variants)(Figure 5C). The extrapolated values for the PLP dissociation rates also showed faster dissociation of PLP from the complex in the D81 set (2.0±0.3 ·10^−3^ s^−1^) to that found for S81 and A81 sets (3.7±1.4 ·10^−4^ s^−1^)(Figure 5D). Overall, the D81 set bound PLP with 70-fold lower affinity than sets S81 and A81 (317±71 μM vs. 4.5±1.2 μM, Figure 5E), which implies a binding energy penalization of about 2.5 kcal·mol^−1^. Thus, the presence of p.S81D affects both the unbound state of AGT as well as its complex with PLP, slowing down PLP binding (penalizing with ~1.5 kcal·mol^−1^) and accelerating dissociation of bound PLP (penalizing with about ~1 kcal·mol^−1^). We must note that the *K*_d_ value derived for AGT WT is one order of magnitude lower than those determined by equilibrium titrations, possibly due to higher uncertainty in the determination of *k*_off_ (as previously shown by [25]). Nevertheless, our data are well consistent with the p.S81D mutation largely slowing down PLP binding and this seems to be the origin of a lower binding affinity.

**Figure 5.**
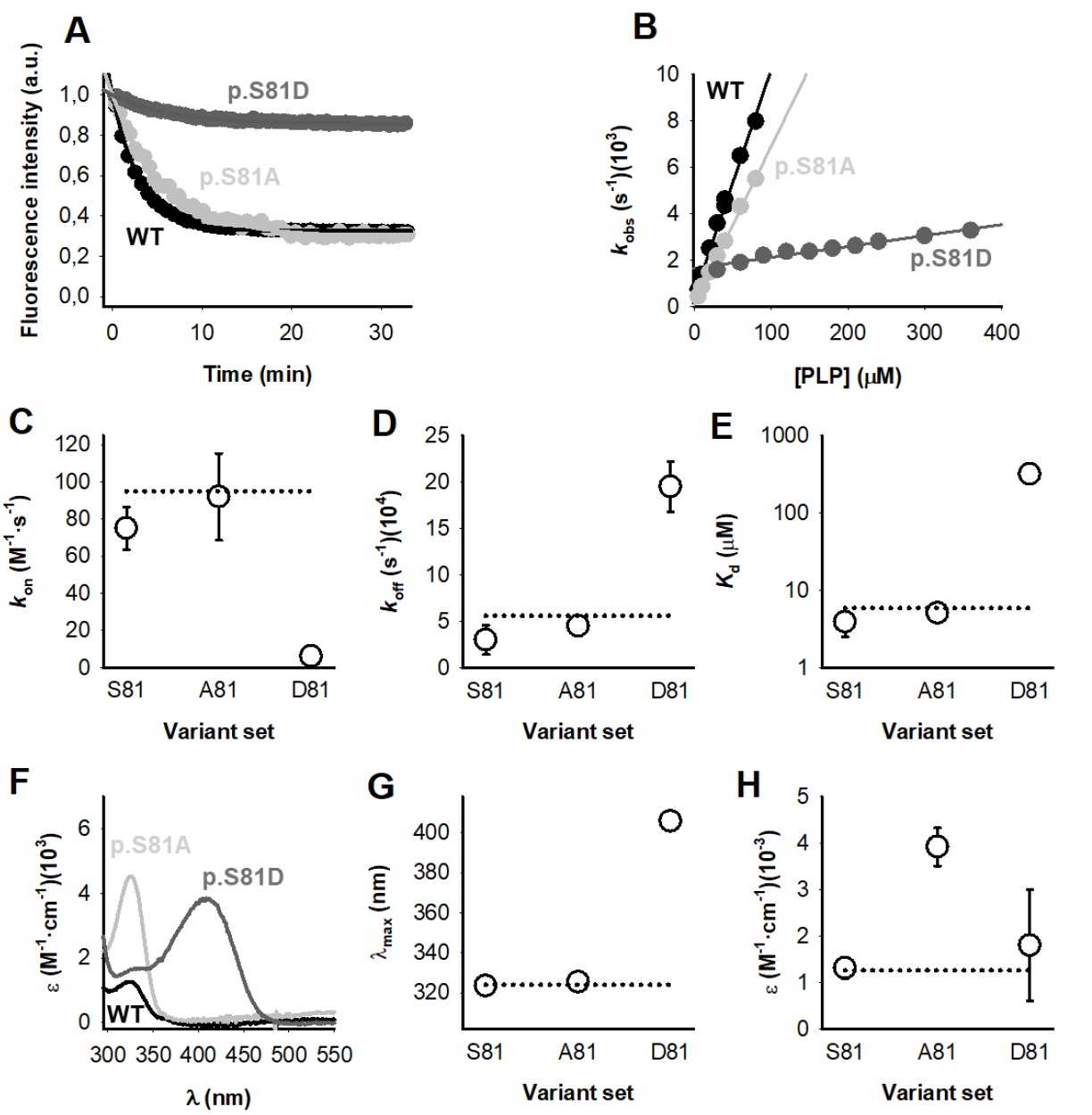
The mutation p.S81D dramatically alters PLP binding affinity and kinetics whereas the mutation p.S81A affects the stability of the PMP-bound state. A) PLP binding kinetic traces (80 µM PLP and 0.4 µM apo-AGT) monitored by fluorescence spectroscopy (Exc. 280 nm, Em. 340 nm). Lines are fittings to a single exponential function which provide *k*_obs_. B) Plots of *k*_obs_ vs. [PLP]. Lines are fittings to a linear function which provide values for the *k*_on_, *k*_off_ and *K*_d_ (Equations 1 and 2).C-E) Values for *k*_on_ (C), *k*_off_ (D) and *K*_d_ (E) for AGT variants belonging to S81, A81 and D81 sets. Note the logarithmic scale in panel E. F-H) Absorption spectroscopic analyses of the filtrates of reaction mixtures containing 20 µM holo-AGT + 200 mM L-Alanine. Panel F shows representative average spectra (for two different preparations of each AGT variants). Panels G and H show values of the maxima of absorption specta (G) and the extinction coefficient at the maximum (H) for filtrates. Data in panels C-E and G-H are mean±s.d. of four different variants in the S81, A81 and D81 sets.

Additional spectroscopic analyses after the first half of the transamination cycle showed that PLP bound to those variants containing p.S81D is readily released from AGT-PLP complex (Figure 4C). Interestingly, we did also observe some effect of the p.S81A mutation on the release of PMP (Figure 4C). Although the maximum of the PMP released after incubation of AGT variants with L-alanine is not affected by p.S81A (324±1 vs 326±1 nm, variants containing in the S81 vs. A81 set), the amount of PMP released from these complexes seems 3-fold higher for the variants containing p.S81A. Considering a ε_325_ for PMP of 8300 M^−1^·cm^−1^ [37], a fraction of PMP released per AGT monomer of 0.47±0.06 is calculated for those variants containing p.S81A, while this value dropped to 0.16±0.03 for those containing S81.

### 3.4. Modeling of the p.S81D and p.S81D mutants structurally support their effects on PLP and PMP binding and catalysis

To characterize the effects caused by p.S81D and p.S81A on the binding of PLP and PMP, as well as their consequences on catalysis, we investigated the AGT-PLP complex (PDB ID: 1H0C) and modeled an AGT-PMP complex, including changes in S81 to A81 or D81. PLP is covalently bound to K209 and thus it cannot move as freely as PMP in the active site of AGT. As a result, the mutation p.S81A does not cause a large perturbation, although the PLP slightly pivots towards W108, a displacement that ultimately accommodates by a coupled movement between H83 side chain and PLP which helps maximizing the interaction between both of them despite the mutation (Figure 6A and C). However, the mutation p.S81D introduces a negative charge near the phosphate group of PLP molecule promoting a larger conformational change that moves PLP further apart from H83 preventing it to establish a hydrogen bond with PLP ligand thus leading to a loss of interactions between the ligand and the enzyme (Figure 6A and D).

**Figure 6.**
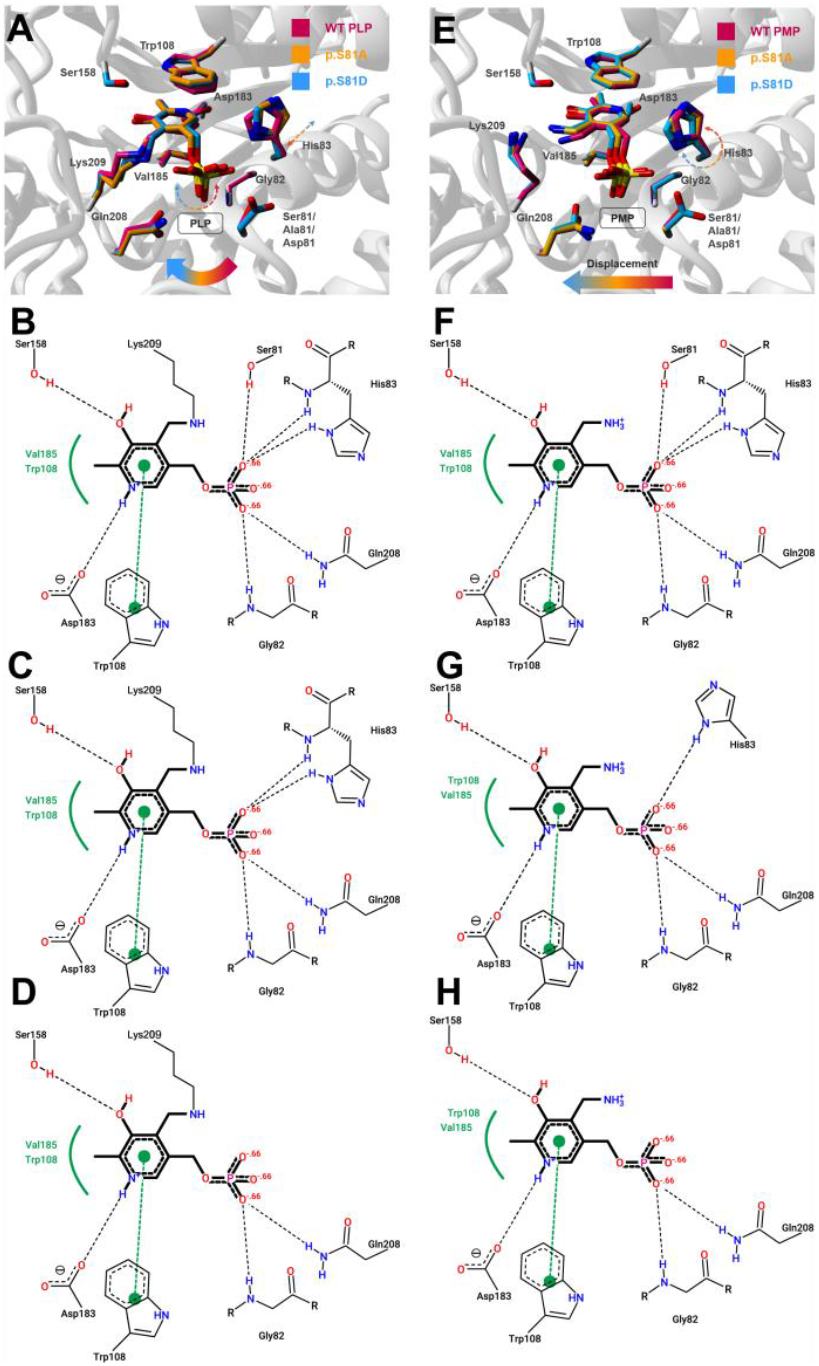
Modeling of the interactions established between PLP (Panels A-D) and PMP (Panels E-H). A-D) Zoom at the active site of AGT-PLP complex (A). PLP and side chains interacting with are displayed as sticks (the color code for WT and the p.S81A and p.S81D mutants is shown). The 2D interaction diagrams of PLP with its neighboring residues at the AGT active site (B, WT; C, p.S81A and D, p.S81D). E-H) Zoom at the active site of AGT-PMP complex (E). PMP and side chains interacting with are displayed as sticks (the color code for WT and the p.S81A and p.S81D mutants is shown). The 2D interaction diagrams of PLP with its neighboring residues at the AGT active site (F, WT; G, p.S81A and H, p.S81D). In panels A and E thin arrows are shown to highlight perturbations upon mutation. In the 2D diagrams H-bonds are shown as black dashed lines, and the hydrophobic contacts are represented by residue labels and spline segments along the contacting hydrophobic ligand parts.

In the AGT-PMP complex, S81 forms a hydrogen bond with phosphate group from PMP, as observed for the PLP in the crystal structure (Figure 1B and Figure 6 B and F). This interaction may hold PMP in place within the enzyme active site and positions it in an optimal conformation. This interaction could play a relevant role as far as can be seen when the mutation p.S81A is introduced. Upon loss of this hydrogen bond, the PMP gains freedom of movement which further weakens hydrogen bonds with H83 (Figure 6 E and G). This results in changes in the binding pose of PMP which involves flipping by approximately ~21º away from H83 explaining the loss of interaction with this residue. The mutation p.S81D disrupts hydrogen bonds with H83 as it pushes the PMP even further from it and flips it by ~37º. As a consequence, less hydrogen bonds are formed between PMP and the enzyme active site (Figure 6 E and H). D81 in turn brings H83 side-chain closer (by getting flipped around the C_α_-C_β_ bond by approximately 14º) in order to establish a hydrogen bond with it which can help stabilizing the N-terminus of the helix α5 where H83 is located

### 3.5. AGT protein noirmalylize with the peroxisomal marker PMP70 both in constructs carrying S81 and those with D81

AGT mostlty targets to the peroxisomes where it is metabolically useful [20,21,38]. To test the possibility of mutation 821D might affect the intracellular location of AGT, we carried out immunofluoresce colocation studies in CHO cells transfected with different variants (WT, LM, WT-p.S82D and LM-p.S82D). Peroxisomal location was large and no mitochondrial location was observed (Figure 7).

**Figure 7.**
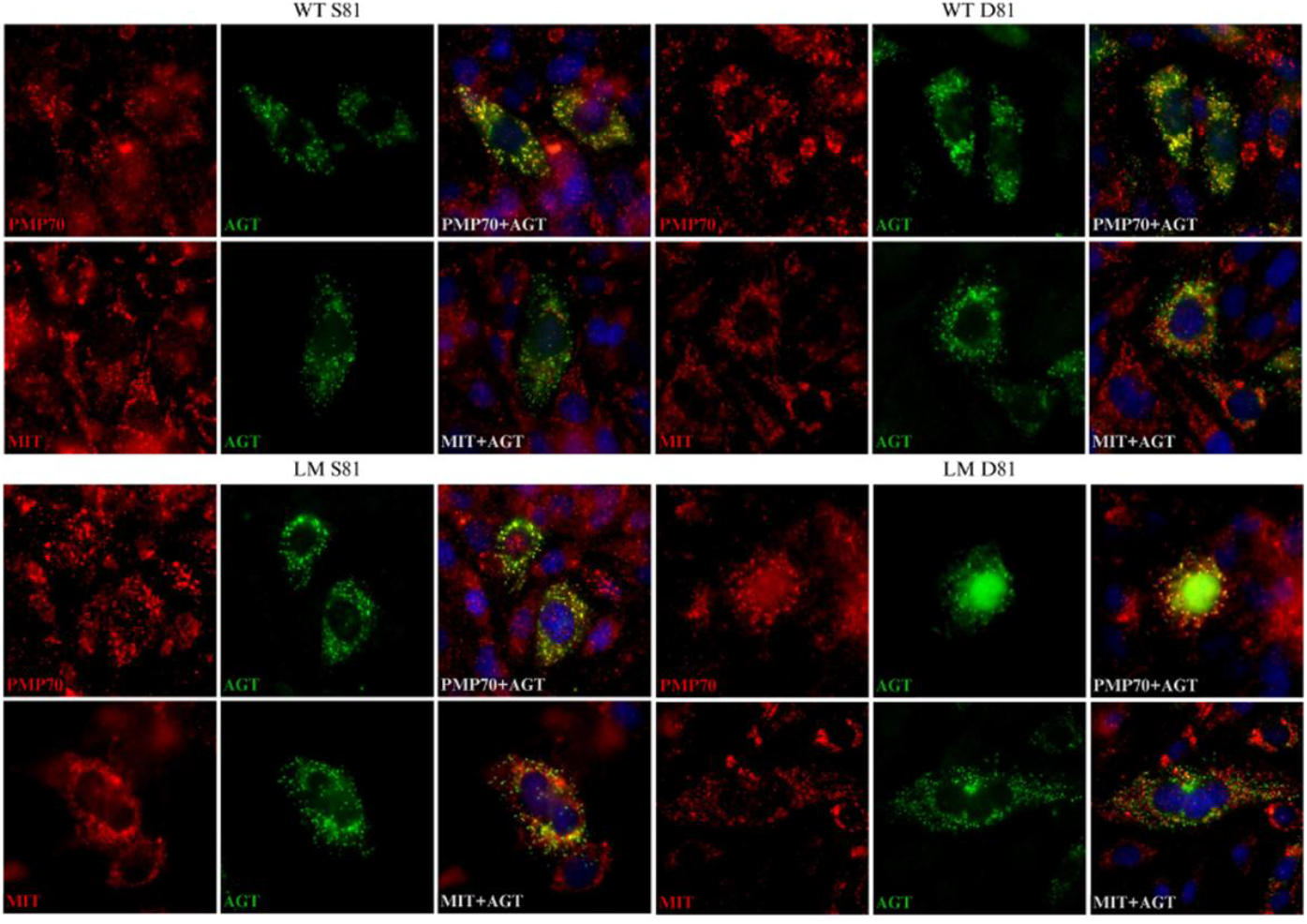
The phospshomimemicc mutation does not affect peroxisomal location. AGT is shown in green while either peroxisomes or mitochondria are labeled in red. All four AGT variants showed peroxisomal localization (colocalizing with PMP70), with the merged images showing yellow signal. No mitochondrial mistargeting is observed, as the localization of the expressed AGT protein within the cell is different from the mitochondrial labeling and the pattern is similar between the WT and LM samples.

## Conclusions

Our structure-function studies revealed that phosphorylation at S81 may reversibly inactivate AGT, depending on those signals controlling phosphorylation/dephosphorylation dynamics (currently, kinases/phosphatases involved are unknown). Phosphorylation of S81 may switch on/ff AGT activity while resididing in peroximes. It may be an additional factor to be considered for genotype-phenotype correlations in PH1. S81 may play an important role in the stabilization of the interaction between the enzyme and PLP/PMP molecules, being more important in the latter. The introduction of a net negative charge disrupts the net of interactions in which phosphate group from PLP/PMP takes part together with the enzyme active site and it also hinders the correct positioning of both PLP and PMP molecules, leading to a decrease and/or abrogation in AGT activity. Interstingly, our work with the mutation p.S81D show functional, structural and electrostatic changes that resemble well those described for PH1-causing mutants such as p.G82E and p.H83R [23,25].

---

**Table 1.** PLP binding to different AGT variants.

|  | **K<sub>a</sub> (M<sup>-1</sup>)(10<sup>5</sup>)** | **K<sub>d</sub> (μM)** | **k<sub>on</sub> (M<sup>-1</sup>·s<sup>-1</sup>)** | **k<sub>off</sub> (10<sup>4</sup>)(s<sup>-1</sup>)** |
| --- | --- | --- | --- | --- |
| **WT** | 1.7±0.4 | 5.9±1.4 | 95±3 | 5.6±1.2 |
| **p.S81A** | 2.4±0.6 | 4.2±1.1 | 132±3 | 5.4±1.2 |
| **p.S81D** | 0.029±0.003 | 345±35 | 4.7±0.3 | 17±1 |
| **LM** | 4.5±1.2 | 2.2±0.6 | 68±1 | 1.5±0.4 |
| **LM-p.S81A** | 1.7±0.3 | 6.0±1.1 | 79±2 | 4.7±0.8 |
| **LM-p.S81D** | 0.038±0.007 | 263±30 | 6.1±0.4 | 18±2 |
| **LM-p.G170R** | 2.4±0.5 | 4.2±0.9 | 70±1 | 2.6±0.7 |
| **LM-p.G170R/p.S81A** | 2.2±0.3 | 4.7±0.6 | 83±1 | 3.9±0.4 |
| **LM-p.G170R/p.S81D** | 0.043±0.012 | 240±70 | 8.0±1.5 | 19±2 |
| **LM-p.I244T** | 3.0±1.9 | 3.3±2.1 | 67±3 | 2.2±1.5 |
| **LM-p.I244T/p.S81A** | 1.9±0.6 | 5.3±1.7 | 74±2 | 3.8±1.1 |
| **LM-p.I244T/p.S81D** | 0.024±0.007 | 421±119 | 5.7±1.4 | 24±4 |

## Supplementary Information

**Figure S1.**
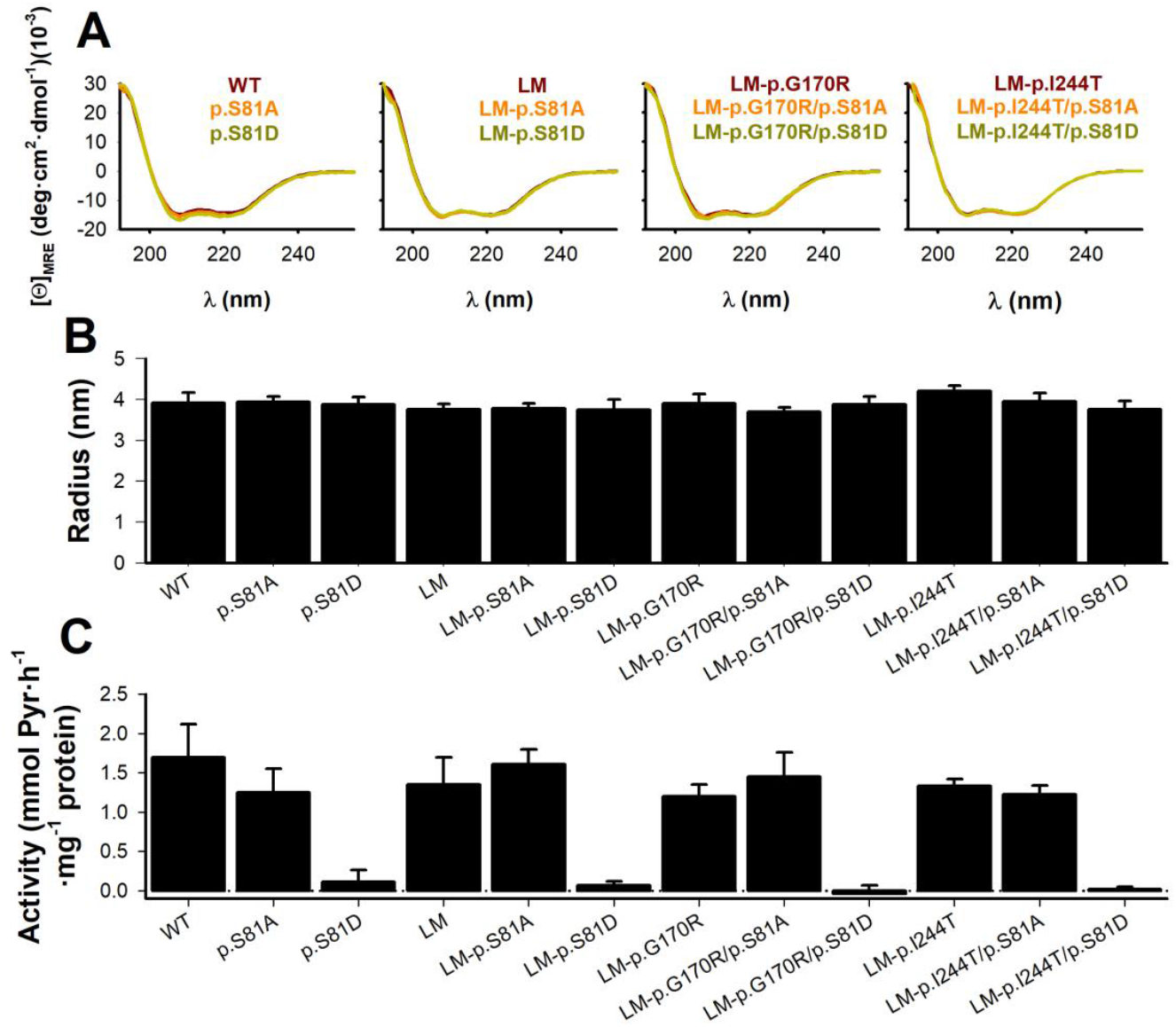
Conformation and specific activity of individual AGT proteins. A) Far-UV CD AGT proteins as purified. For each variant, spectra are the average from two different purifications. B) Hydrodynamic radius for AGT proteins as purified from DLS analyses. Data are the average ± s.d. from 4-6 independent measurements. C) Overall transaminase activity of AGT variants using L-Alanine (100 mM) and glyoxylate (10 mM) as substrates. Data are the average ± s.d from at least four experiments.

**Figure S2.**
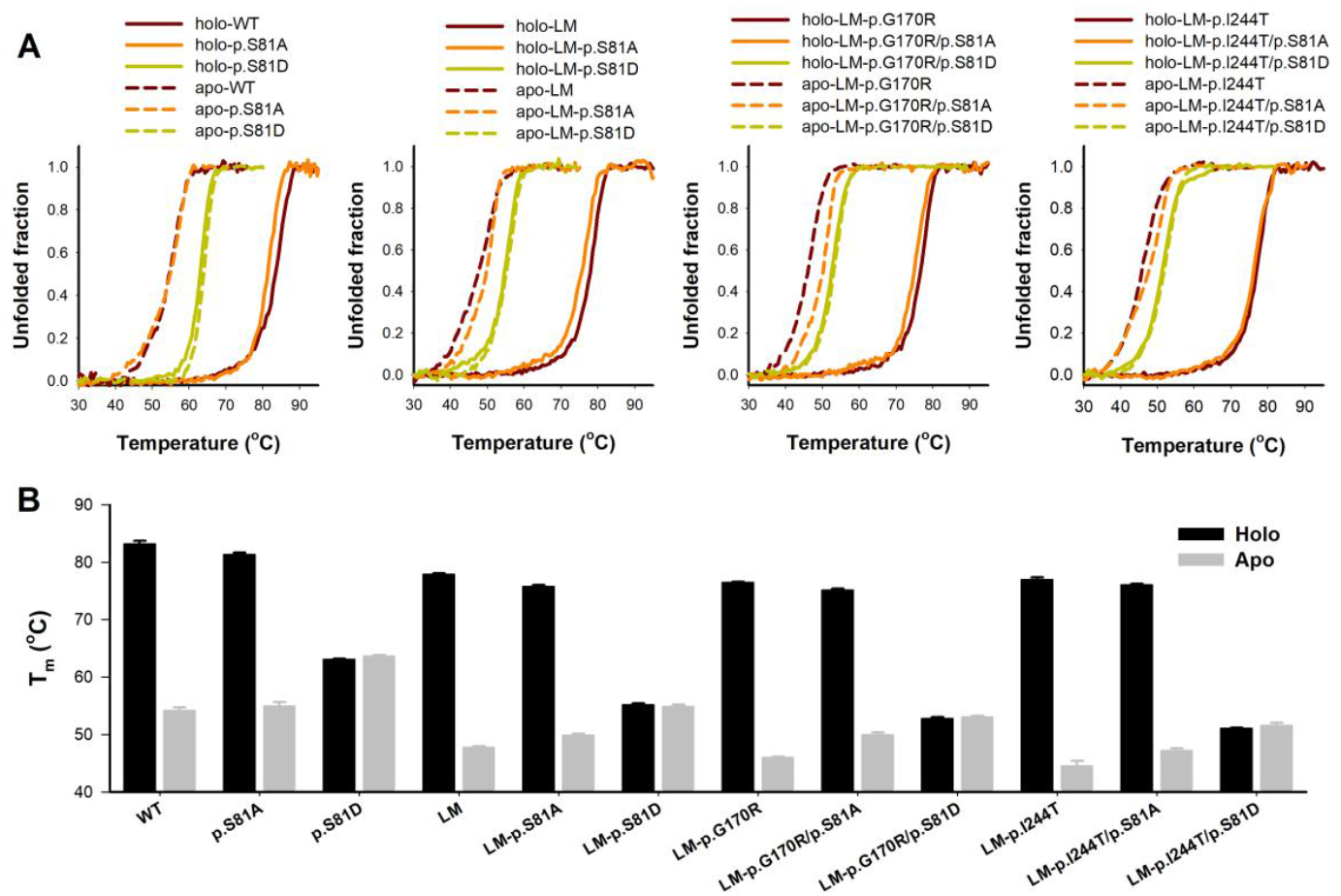
Thermal denaturation of holo- and apo-AGT individual variants. A) Representative thermal scans. B) T_m_ values (mean±s.d. from four replicas).

**Figure S3.**
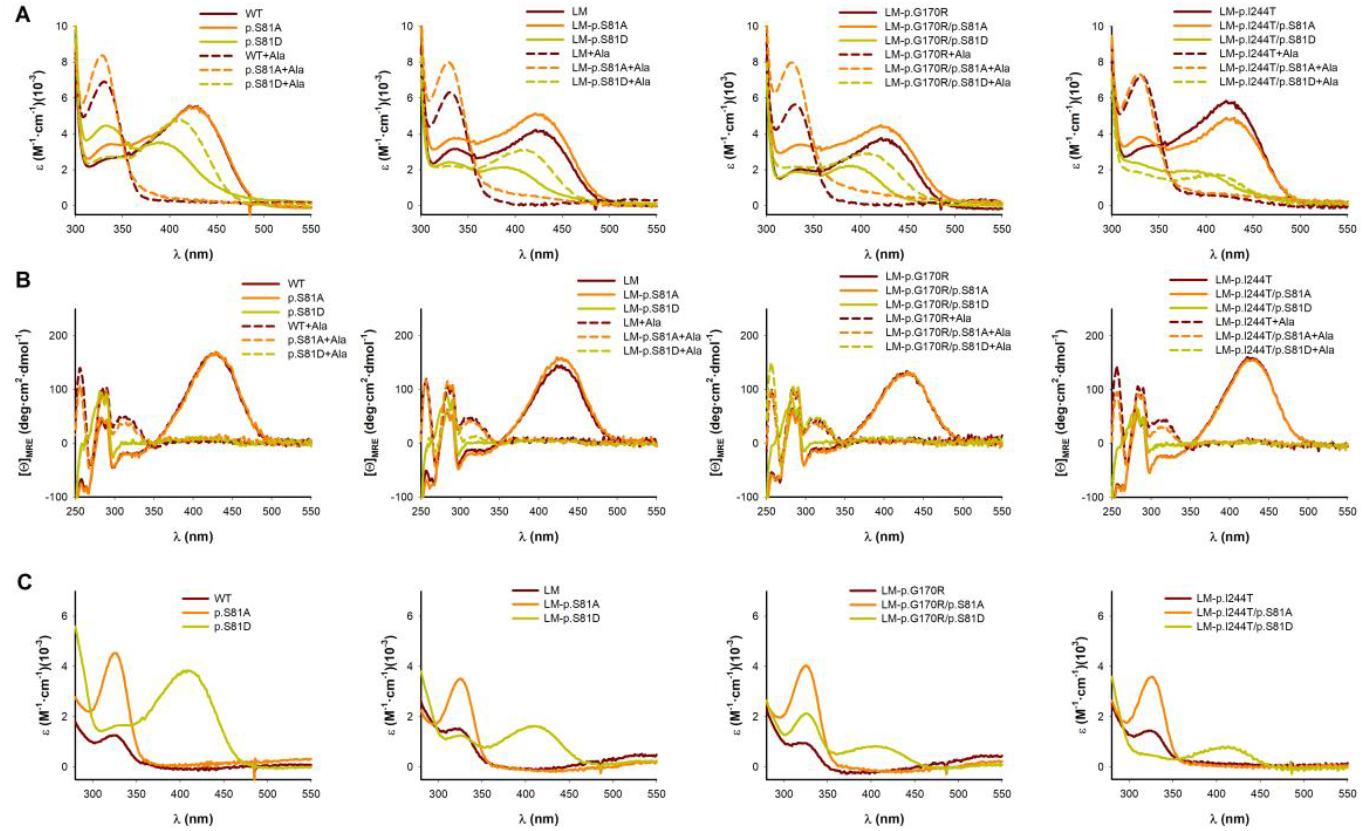
Spectroscopic characterization of PLP and PMP along the functional cycle of AGT variants. A-B) Absorption (A) and CD (B) spectra of AGT proteins (15 μM) with and without incubation with alanine 200 mM. C) Absoption spectra of reaction mixtures containing AGT and alanine 200 mM (panel B) filtered using a 30 kDa concentration devices.

## Notes

### Competing Interest Statement

The authors have declared no competing interest.

### Funding

- ERDF/Spanish Ministry of Science, Innovation and Universities-State Research Agency: Grant number RTI2018-096246-B-I00

